# Biomechanics of stem cell fate decisions in multilayered tissues

**DOI:** 10.1101/2024.12.12.628138

**Authors:** Preeti Sahu, Sara Canato, Raquel Soares, Adriana Sánchez-Danés, Edouard Hannezo

## Abstract

Tissue homeostasis relies on a precise balance of fate choices between renewal and differentiation, which is dysregulated during tumor initiation. Although much progress has been done over recent years to characterize the dynamics of cellular fate choices at the single cell level, their underlying mechanistic basis often remains unclear. In particular, although physical forces are increasingly characterized as regulators of cell behaviors, a unifying description of how global tissue mechanics interplays with local cellular fate choices is missing. Concentrating on skin epidermis as a paradigm for multilayered tissues with complex fate choices, we develop a 3D vertex-based model with proliferation restrained in the basal layer, showing that mechanics and competition for space naturally gives rise to homeostasis and neutral drift dynamics that are seen experimentally. We then explore the effect of introducing mechanical inhomogeneities, whereby subpopulations have differential tensions. We uncover that relatively small mechanical disparities can be sufficient to heavily tilt cellular towards symmetric renewal and exponential growth. Importantly, the simulations predict that such mechanical inhomogeneities are reflected by distinct morphological changes in single-cell shapes. This led us to derive a master relationship between two very different experimentally measurable parameters, cell shape and long-term clonal dynamics, which we validated using a model of basal cell carcinoma (BCC) consisting in clonal Smoothened overexpression in mouse tail epidermis. Altogether, we propose a theoretical framework to link mechanical forces, quantitative cellular morphologies and cellular fate outcomes in complex tissues.

## Introduction

Epithelial tissues have a central barrier function in multicellular organisms, and undergo extensive turnover during homeostasis. The skin epithelium, the largest organ in our body, renews itself completely within four weeks for example [18,29].This crucial process of homeostatic self-renewal relies on the basal proliferative layer of the interfollicular epidermis. This layer is composed of stem cells and progenitors, which divide and give rise to differentiated suprabasal cells, showing balanced cell fate during homoestasis and unbalanced behaviours that can lead to wound healing or cancer initiation [2, 3, 6, 12, 13, 24, 27, 41, 44, 55]. Although lineage tracing datasets have proved instrumental over the last 15 years in defining the potential, hierarchy and modes of divisions of different cellular populations, key open questions remain as to the mechanisms underlying fate choices. For instance, in the context of the homeostatic skin epithelium which follows long-term neutral drift due to stochastic basal cell renewal, what are the mechanisms which determine whether a given basal cell division will give rise to two basal (or suprabasal) cells, i.e. a symmetric division, or one basal and one suprabasal cell, i.e. an asymetric division? Furthermore, at homeostasis, these processes have to be balanced at the population level to ensure a conserved cell number, and this balance is lost in a predictable manner in processes such as wound healing [12, 44] or basal cell carcinoma initiation [2, 55]. Based on previous works, determinants of cell fate decisions can be grouped into two generic categories– (a) cell-intrinsic i.e. the underlying gene regulatory networks [34] and (b) cell-extrinsic factors i.e. local *niche* specific bio-chemical cues [9, 10] or biochemical interactions between different stem cells [19, 26]. However, from a theoretical perspective, all of these types of models give rise to very similar long-term behavior. For instance, both stochastic zero-dimensional models (where cellular decisions are cell-intrinsic and do not consider space) or stochastic Voter models on spatial grids (where a cell can differentiate/move suprabasally upon the symmetric division of neighboring cells) give rise to identical clone size distribution and long-term patterns of clonal growth [19, 23–25, 41, 44].

While the predictive power of such minimal models has been impressive, the next emerging question is thus to understand more mechanistically how the fate decision probabilities that underlie these stochastic models are encoded. In spermatogenesis, it has been recently shown that competition for diffusible fate determinants is a simple homeostatic mechanism able to collectively balance cellular fate decisions, which can explain a number of homeostatic and perturbed conditions [21]. Furthermore, the impact of cellular forces and mechanics has been increasingly recognized over the last decade. Since the classical observation that substrate rigidity can modulate single cell fate decisions in vitro [36], a number of emerging evidence also suggest impact for cell and substrate mechanics in tissues [1, 16, 32, 44, 45, 58]. However, the impact of mechanical forces on 3D cellular fate choices remain poorly explored from a theoretical perspective, which we tackle in this work.

Over the past decade, extensive efforts in the field of mechanobiology have been devoted to developing minimal theoretical descriptions of 2D and 3D tissue mechanics. One of the simplest and most successful description of confluent tissues has been vertex and voronoi models models, where cells are represented by polygons with tensions associated to their shared cell-cell contacts [4, 8, 15, 17, 30, 35, 53]. This is based on the fact that most cellular forces are concentrated at interfaces, from the cell cortex and adhesion machinery [15, 39]. The equilibrium shape of cells and tissues can then be derived by minimizing the total mechanical energy of the system, and can predict a number of complex collective behavior such as density-independent unjamming transition, novel mechanisms for cell sorting or shear-thickening [4, 5, 28, 30, 47–50, 54] many of which have been observed in biological tissues [14, 40, 51, 56, 57]. However, these models are traditionally been confined to describing tissues in the absence of cellular renewal or fate choices, e.g. tissues made of a single cell type, or of two independent populations undergoing sorting. To develop a unified theoretical description taking into account both tissue mechanics and cellular fate choices, we choose to concentrate on the skin interfollicular epidermis (IFE), which comprises of several confluent layers of cells in homeostasis. Only cells localised to the bottom, or basal, layer (in contact with the basement membrane, or BM) can divide in this system, while suprabasal cells are lost at a constant rate, creating a steady flow of cells upwards. As a first approximation and based on the “open-niche” nature of the skin epithelium [41], we assume that all basal cells are identical.

We first show that a ‘balanced’ mode of proliferation emerges organically from simply allowing the SCs to make mechanically favourable fate choices i.e. in absence of any spacial bias or pre-assignment of fates. We then test how fate choices can be modulated by inhomogeneities in the mechanical properties of somes cells, in particular differences in basal adhesion to the BM. We find that clones with lower basal tension undergo a distinct change in their proliferation mode, due to the fact that remaining basal after division becomes mechanically favorable. Interestingly, fate outcomes are highly sensitive to mechanical inhomogeneities, with 10% decrease in tensions being sufficient to transform clones into nearly exponential growth mode. Furthermore, we show that the amount of tension inhomogeneity can be extracted based on cell shape alone, providing a key relationship between cell shape and cell fate outcomes - two experimentally accessible parameters. We test these in-silico predictions in mouse tail interfollicular epidermis (IFE), where oncogenic Smoothened activation has been shown to drive BCC initiation due to a well-defined tilt of fate outcomes towards symmetric renewal. We find a small but significant change in cellular shape upon oncogenic activation, which can predict the range of clonal fate imbalance observed in the data.

## Model

We start by using the vertex model, a classical framework to study two and three dimensional tissues. The mechanical energy of the tissue is a sum of its contribution from the surface (BM) and the bulk (individual cells). Each cell is associated to an energy, written as [30]

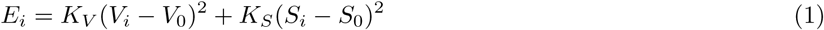

where *S*_0_ is the preferred surface area of each cell, *V*_0_ its preferred volume, and *K_S_* and *K_V_* the compressibilities associated to how strong these regulation towards these preferred area and volume are. The preferred cell shape index, defined as 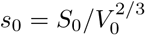, is a simplest representation of the different types of forces acting at the interface of different cells, and has been shown to be a simple predictor of the collective physical state of the tissue, with *s*_0_ *>* 5.41 giving rise to fluid tissues (where cell-cell re-arrangements cost vanishing energy) and *s*_0_ *<* 5.41 to solid tissues (where energy barriers associated to cell-cell re-arrangements result in tissues able to sustain mechanical load). Here we use a solid-like value of *s*_0_ = 5.30 for all the cells. Given this homogeneity, we can use *K_V_* = 1 [50]. Preferred volume is set as *V*_0_ = 1, thereby setting the length scale of the simulation. We build an in-silico epithelial tissue composed of three layers of cells, as shown in Fig1a-c, where each cell is a polyhedral unit created using the Voronoi tesallation of all the cell centers-which are the degrees of freedom of this system [49]. Random noise is added at each time point to the cell centers, as sketched in Fig1a to explore the effect of random motility and/or mechanical fluctuations in the system, which has been shown to be an alternative route to tissue fluidization [5, 20, 33, 42] compared to changes in shape index.

**Figure 1.**
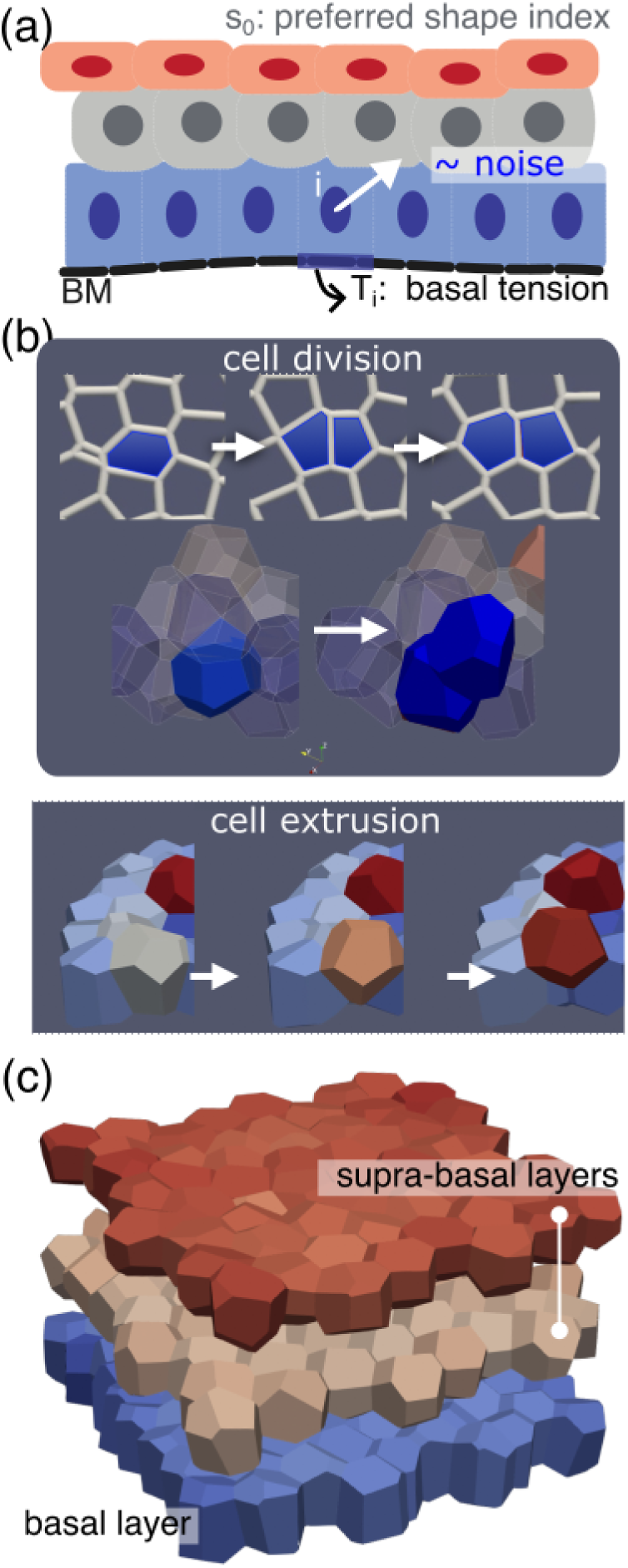
Schematic of the model and snapshot of simulations. (a) Schematic of our model for multilayered tissues on top of a basement membrane (BM) shown in black. Each cell has a preferred cell volume *V*_0_(*t*) and cell shape *s*_0_. The color scheme denotes the distance of the cell from the BM – the closest to the furthest cells are colored from blue to red. Noise is added to the cell center (blue sphere). Cell-BM contacts have an additional interfacial tension *T_i_*, rending the interface relatively flat. (b) Cell division event is depicted first in the the basal cross-sectional view and below in 3D space: the randomly selected cell (in blue) divides into two daughter cells, that gradually grow back to the original volume. Cell extrusion event occur organically, defined as a basal cell (in blue) being pushed out towards the suprabasal layer (in red), i.e. shrinking its basal area to zero. (c) Tissue is shown in its steady state (after one complete turnover time of *τ_D_*). The tissue dimensions are 8*x*8*x*3(*unit length*)^3^ and contains a total of 192 cells that are divided into basal layer (blue) and the supra-basal layers above (red + orange). Boundary conditions are periodic along the horizontal x and y axes.

Next, we sought to model the interface between the epithelium and the basement membrane. To do this in a minimal manner, we assume a high tension at the basal-BM interface, which results in almost flat BM. Note that we do not explicitly model the elastic properties of the BM given that it remains largely undeformed during the early processes that we will study, and that we use values of interface tension intermediate enough to reduce artificial pinning between basal cells and the medium below (see Sec:Methods Text for more details and sensitivity analysis) [49, 54].

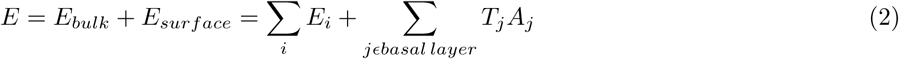

Finally, in IFE, as in other stratified tissues, the bottom-most cellular layer (composed of basal cells) adheres to a rigid basement membrane (BM), as sketched in 1a, and can proliferate, whereas layers above (suprabasal cells) cannot divide and are progressively shed as they undergo differentiation. Thus, in simulations, we allow basal cells i.e. any cell with a sharing in contact with the BM, to grow in size and divide at a random rate *k_D_* (see Sec:Methods text for more details). The growth of mother cell is implemented by gradually increasing the preferred volume-*V*_0_. After this occurs, we select a random plane of division (perpendicular to the BM, shown in Fig1b) and add another cellular center. To conserve the total number of cells, at each division, we also randomly pick a suprabasal cell for deletion. This means that we do not impose any cellular fate balance in basal cells, and each cell can transit from the basal to the suprabasal compartment organically as a result of force balance as shown in Fig1b.

### In-silico tissues follow balanced neutral-drift dynamics

Importantly, we find that simulating this minimal system organically displays a spatial and dynamical organization close to their in-vivo counterpart. Firstly, the basal layer displays apico-basal oriented prismatic shapes whereas suprabasal cells are more isotropic (as visible in Fig1c and quantified in Fig S1 a-b), which comes from the constraint with the basement membrane (similar to previous observation at the boundary of two different tissues [49]). However, because of low tension values, there is negligible crowding due to basal tension (details in Fig S1 c). Secondly, the system organizes itself into a state of permanent flux from basal to suprabasal layer. This is because cells staying in the basal layer after division would result in crowding, which is energetically unfavorable compared to moving to suprabasal layers. However, basal proliferation still result in an increased basal density, as quantified in Fig S1 c. Thirdly, given that all basal cells are modelled as identical, the ones mechanically poised to extrude towards the suprabasal layers is a random process, resulting in neutral drift dynamics within the epithelium. To visualize this and measure it more quantitatively, we performed computational lineage tracing in our simulations, assigning to each cell a given ID that is retained upon division. Given the difficulty of long-term live imaging in this system, lineage tracing has been a key experimental method to assess cellular fate choices, by irreversibly labelling sparse populations of cells as well as all their future progeny (clones), and measuring clone size at given time points post labelling [55].

In accordance with previous analytical results, we found that as cells divided and extruded from the basal layer, the number of surviving clones declined at 1*/t* as the average surviving clone size grew linearly (Fig2a-c). Reconstructing lineages was consistent with this picture, with around 25% of basal cell divisions resulting in two basal cells, 50% in one basal and one suprabasal cells and 25% in two suprabasal cells. These are the results one would expect if following a division, the two daughter cells have an uncorrelated 50-50 probability to extrude suprabasally. Interestingly, this is quite close to values observed in short-term live imaging in mouse ear epidermis [43].

**Figure 2.**
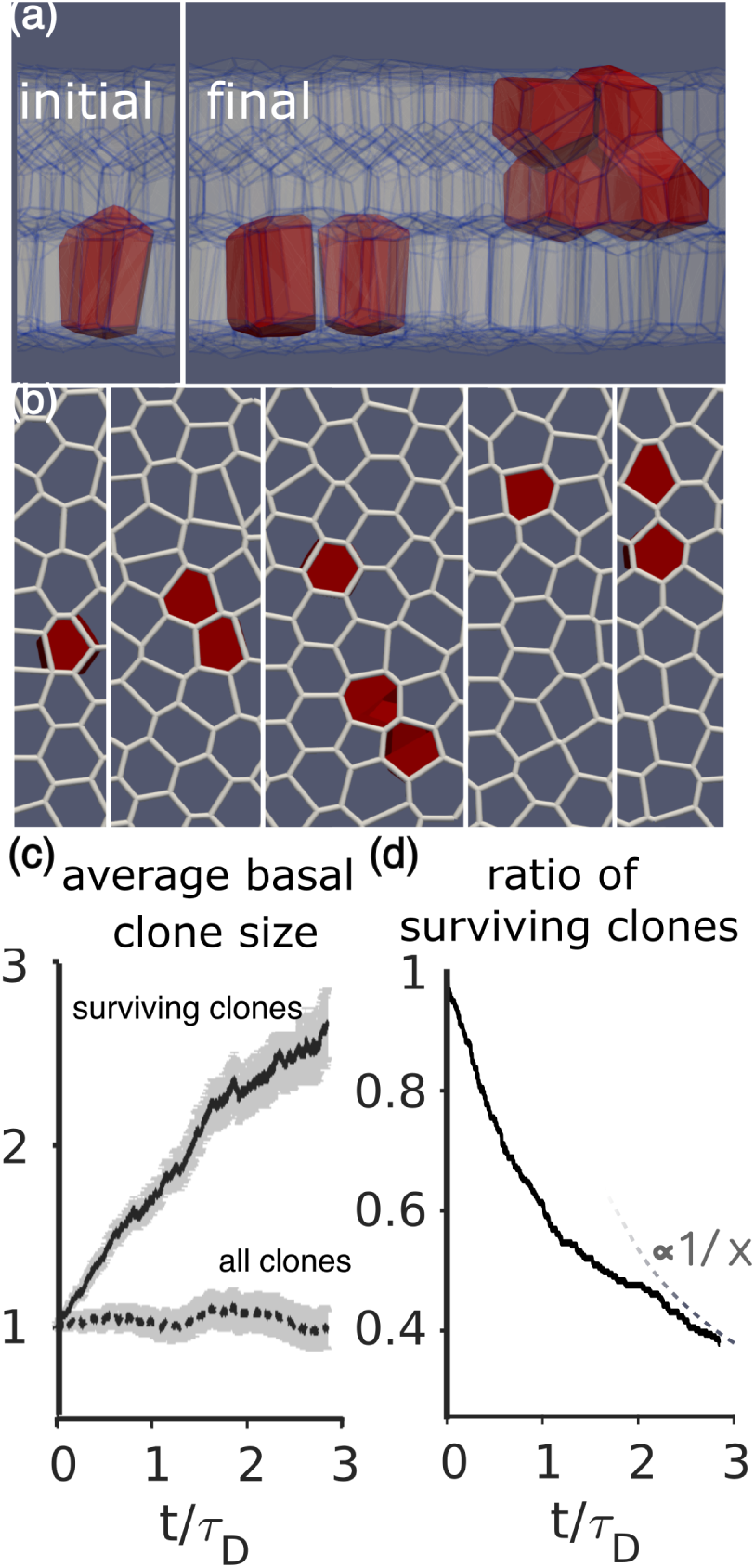
Homogeneous mechanical environment displays neutral drift dynamics. (a-b) Clonal growth dynamics of a representative clone is depicted in 3D (a) and in the 2D basal plane (b). Cells that belong to the same clone are tagged in red. Almost three turnovers later i.e. despite several divisions, the number of basal cells has only doubled, while the rest have extruded to suprabasal layers– as shown in the final frame. The same clone is shown in the 2D basal cross-section (b), where in addition to the initial and final frames, one-forth, middle and three-fourth frames are also shown. (c) The average number of basal cells is plotted with respect to time (t) in units of turnover/division timescale *τ_D_*. When averaged over all the clones, the size remains almost constant at unity– only to be expected as the total number of cells is conserved. Averaging over the surviving clones, it shows an initial transience, followed by linear growth– as seen in neutral-drift processes. (d) The ratio of surviving clones decreases inversely with time, in line with the above quantification. Parameter set: Total number of cells = 192, *s*_0_ = 5.3, *T* = 1.0 and zero noise.

Next, we tested if these results were sensitive to the mechanical state of the tissue, by varying the shape index of cells (i.e. simulate fluid vs solid tissues), the basal tension, as well as the amount of noise in the simulations. Interestingly, we found that the results were largely un-affected in terms of cellular fate dynamics and clone size, as shown in Fig S1 c-f. However, the mechanical state of the tissue did have an impact on geometrical features of clones: for both solid and fluid tissues with high noise, the clonal footprint were more elongated and fragmented ( see Fig S1 g-h for solid and fluid systems respectively) which we quantified using the ‘length’ of a clone i.e. the maximum distance between two basal progenies. Such features could be readily compared to experimental lineage-tracing datasets, which have tended to historically concentrate on overall clonal sizes rather than geometric cellular arrangements. At homeostasis, clones in skin epidermis tend to be highly compact and cohesive [55], which would support a solid state for the tissue.

### Driving unbalanced fate choices by differential cellular mechanics

Next, we sought to relax the assumption of identical basal cells to understand processes such as tumor initiation. Lineage tracing combined to targeted oncogenic activation have made it possible to study the dynamics of tumor growth from single tumor cells in an in vivo context. For instance, Smoothened activation in single cells results in formation of preneoplastic lesions (dysplasia) or invasive BCC depending on the cell of origin (progenitor or stem cell) [55]. In previous works, we had shown that the early stages of tumor formation could be well-recapitulated with a stochastic model of cell fate reminiscent of homeostasis, but with a constant imbalance towards symmetric basal divisions [55]. Surprisingly, the main driver of tumor formation was therefore not enhanced basal proliferation, but instead by decreased differentiation. Recent studies have also investigated the role of the mechanics of the BM and stroma on later stages of tumor invasion [16]. However, these previous works leave open the key question of what sets the imbalance of cellular fates in tumors. Driven by previous works showing the importance of mechanical forces for determining the mode of cell fate choices [1, 32, 37, 38, 44, 56], we sought to test whether these imbalances could have a mechanical origin.

Given the simplicity of our model, there are only few mechanical parameters that can be different in tumors. As described above, enhanced proliferation is not correlated experimentally to unbalanced fate [55], so we investigated the effect of differences in mechanical tensions. In our model, the choice to differentiate is based on a mechanical competition to finite access to the basal layer, and we therefore reasoned that differences in basal adhesion/tension would have profound impact on cellular fate choices and clonal dynamics.

We therefore simulated a system where a few mutant cells and their progeny were endowed with decreased tension to the BM by a ratio *R* = *T_B_/T_T_ >* 1, where *T_B_* and *T_T_* are the base-line wild-type (WT) and mutant basal tensions respectively as shown in Fig3a, resulting in their preferential adhesion to the BM compared to wild-type cells. Strikingly, very small differences in basal tensions were sufficient to drive profound changes in clone behaviors. Indeed, cellular fate choices became heavily biased towards symmetric divisions, with much larger basal clone sizes (Fig3b). This can be quantified in terms of overall clonal growth dynamics, which became exponential rather than the linear trend observed for homeostasis, with a mere 10% decrease in basal tension (Fig3c) being sufficient to give rise to nearly exclusive symmetric divisions. Moreover, by investigating different values of basal tension differences, we found that the effect was progressive, with a monotonic increase in average basal clone size with tension ratio R (Fig3d).

**Figure 3.**
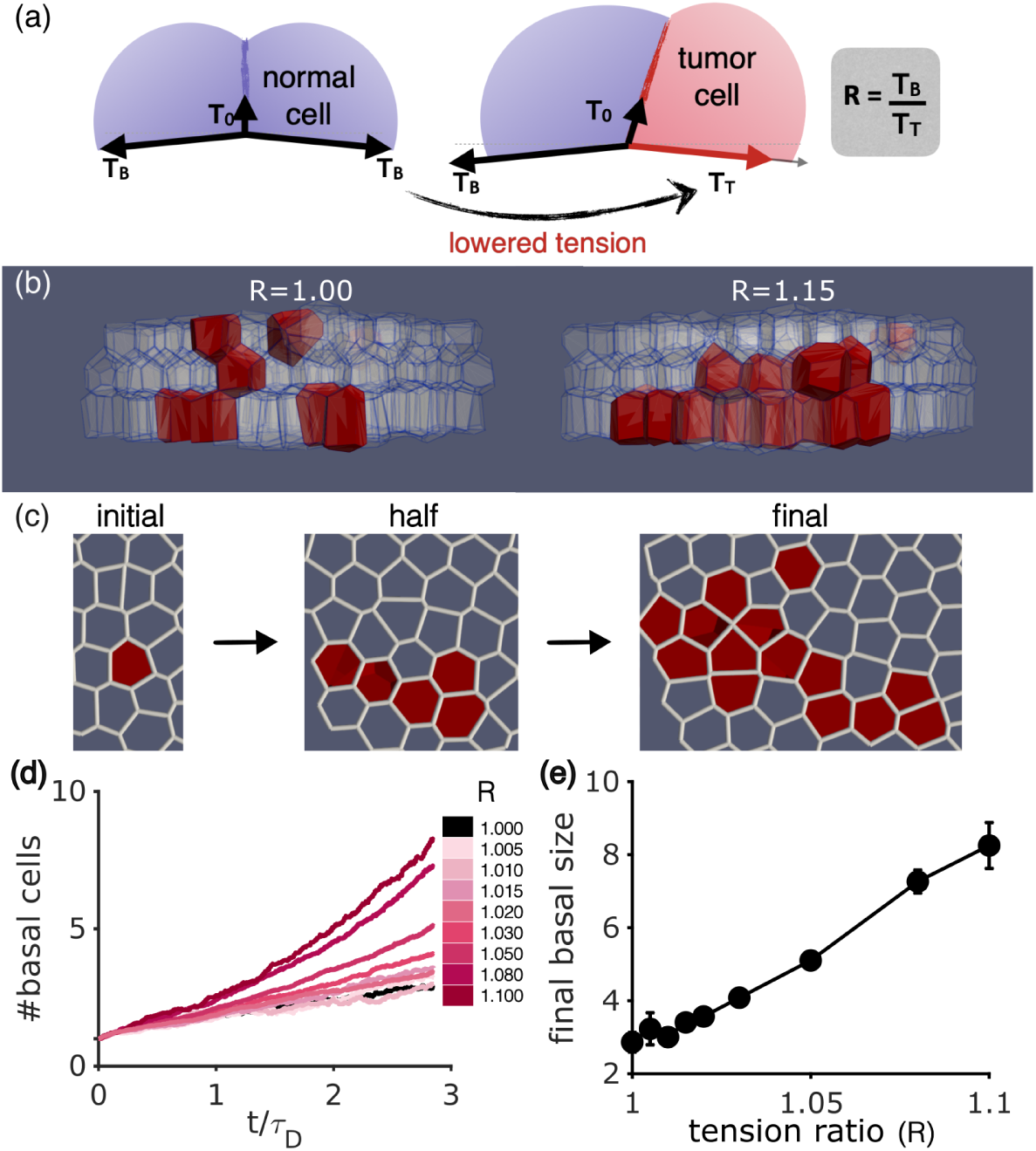
Basal tension changes within a clone induce tumor-like exponential growth. (a) Schematic for introducing tension disparity is shown, wherein normal cells in blue have basal tension *T_B_*, and mutant cells in pink (and their progenies) have a lowered basal tension i.e. *T_T_ < T_B_*, making the ratio *R* = *T_B_/T_T_* ≥ 1. Inter-cellular/bulk tensions are represented as *T*_0_, and are left unchanged. (b) Final 3D morphology is shown for a single clone (red cells) with either normal properties (*R* = 1, left) and a mutant clone with decreased basal tension (*R* = 1.15, right). (c) 2D basal cross-section of the same mutant clone, where initial, middle and final frames are shown, depicting a much faster growth as compared to Fig2b. (d) Number of basal cells, averaged over surviving clones and plotted with respect to rescaled time (*t/τ_D_*), undergoes a transition from linear for *R* = 1 to exponential for *R* = 1.1 (lighter to darker shade of pink). (e) The final basal size, plotted against tension changes *R*, shows a monotonically increasing trend. Parameter set: Total number of cells = 192, *s*_0_ = 5.3, base-line/WT tension value *T_B_* = 1.0 and medium noise(*v*_0_ = 0.1).

Again, we tested how these results were affected by the mechanical state of the tissue (Fig S2 a-b), the degree of noise (Fig S3) and the baseline wild-type value of the basal tension, finding that the changes in clonal dynamics with different tension ratio *R* are highly robust (Fig S2 c). In particular, the number of basal cells remains largely unchanged by these model variations, and mainly affected by the basal tension ratio *R*, although as discussed in the homeostatic case, the spatial shape of clones becomes more dispersed with increasing noise, as shown in (Fig S3). We also observe that that the basal growth dynamics depends mainly on the tension ratio R (rather than the absolute difference between both tensions)(Fig S2 c). Overall, these simulations suggest a key role for basal tension in favoring the adhesion of basal cells to the BM in tumors, a wetting-like phenomena which favors them to stay basal after divisions and thus providing a competitive advantage over surrounding wt cells.

To describe more quantitatively how individual fate choices are affected by tension differences between tumor and wild-type cells, we plotted individual clonal lineage to extract the probability of all three fate outcomes (basal-basal, basal-suprabasal, suprabasal), defining population asymmetry-Δ, as the imbalance between symmetric division and symmetric differentiation Fig4a-c. We found that the the asymmetric fate outcome starts from 50% and slowly declines with increasing R. This population is compensated by an increase in symmetric divisions. While both kinds of symmetric fates are around 25% in the homeostatic case (*R* = 1), they gradually deviate such that by *R* = 1.1, the symmetric differentiation plummets to less than 5% and the symmetric divisions increase to over 70% as shown in Fig4d increasing the population asymmetry Δ with respect to tension ratio R(Fig S2 d).

**Figure 4.**
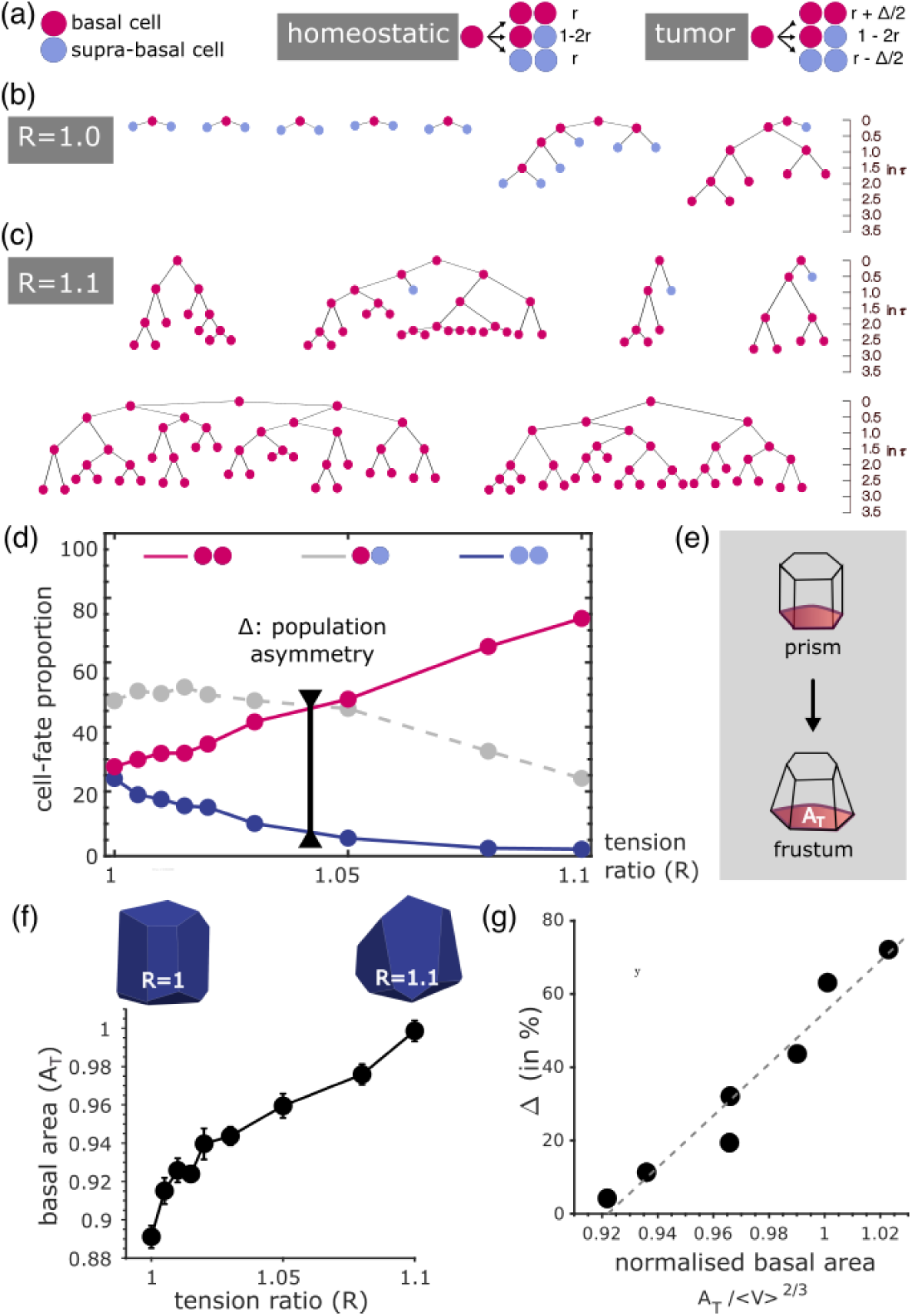
Fate choice imbalance Δ increases as a function tension ratio R, in tandem with cell-shape changes. (a) Schematic shows the probability distribution of homeostatic and tumor fate choices, where Basal and supra-basal cells are depicted in magenta and blue respectively. In balanced mode, the distribution between all three fates (basal-basal, basal-suprabasal, suprabasal) is symmetric, with *r* denoting the basal cell doubling. In tumors, the symmetric fates get imbalanced by Δ. (b,c) Representative examples of lineage trees are shown for tension ratios–*R* = 1.0(above) and *R* = 1.1 (below). Being a basal cell, magenta cells have the power to divide, whereas blue cells typically lose it upon extrusion. All the trees are to scale with time, shown in units of *t/τ_D_* on the right. Magenta cells at the end of a tree have unknown fate as they survive in the basal layer till the simulation ends. The division time corresponds to the appearance of daughter cells, and not the mother cell. (d) Proportion of all three fates– basal-basal in pink, basal-suprabasal in gray, suprabasal in blue, of all cell divisions that took place within the first turnover i.e.*time* ≤ 1, is plotted with respect *R*. The distribution is initially symmetric with values– 25%, 50%, 25%. Δ becomes the difference between magenta and blue curves, that increases with increasing *R* because the proportion of suprabasal cells plummet to zero, and are balanced by the ever-increasing ratio of basal-basal fate. The asymmetric fate shows a moderate decline. (e) Schematic to show the qualitative change in cell shape from prism-like to more pyramidal frustums. Basal area *A_T_* is represented in red. (f)*A_T_* is plotted with respect to *R*. Insets show the representative polyhedral cells that corroborate the increasing trend of *A_T_* with their increasingly wider base. (g) Connection between fate imbalance and cell-shape changes. Fate choice imbalance Δ is plotted with respect to basal area *A_T_* (normalised by the cell volume for the given value of R) in black dashed curve, and shows linear relationship between both.

A strong prediction of our model of mechanically-driven fate choice imbalance is that the shape of tumor cells should be increasingly modified for increasing tension ratio R. Indeed, we verified in the simulations that for increasing tension ratios, tumor cells tended to adopt an increasingly pyramidal shape with expanded basal area, as shown in Fig4e,f - which intuitively reflected their preference for basal adhesion. Importantly, given that this change in cell shape was gradual with increasing *R*, we reasoned that it could be used as an intuitive read-out for tension disparities, which can be readily quantified in experiments. An alternative and complementary metric was the contact angle between lateral areas of the wt-mutant clone interface, which became increasingly tilted for increasing *R* as expected in a simple wetting argument.

Crucially, this means that the model predicts a highly non-trivial, yet robust, correlation between the shape of a tumor cell (a static morphological feature) and the bias in its fate choices between division and differentiation (a complex dynamical property), which is expected to be roughly linear as shown in Fig4g.

### Testing the model in basal cell carcinoma initiation

To test this prediction, we used a genetic mouse model of BCC that consists on the activation of the constitutive form of Smoothened fused to yellow fluorescent protein (SmoM2-YFP) under the control of the Keratin-14 (Krt14-CREER) upon tamoxifen administration, thereafter referred as Krt14-CREER/SmoM2. Administration of low dose of tamoxifen administration leads to the activation of the oncogene in few basal cells of the epidermis in Krt14-CREER mice, leading to well-separated clones. We compared the clones generated upon oncogenic activation to those clones generated in homeostatic conditions by inducing the expression of YFP in the basal cells of the epidermis (Krt14-CREER) upon tamoxifen administration, thereafter referred as Krt14-CREER/YFP. Whole mount stainings followed by confocal imaging and quantification, shows that SmoM2-expression in the basal cells of the epidermis leads to an expansion of the basal compartment, as shown by an increase in the number of basal cells/clone, detected as early as 1 week upon tamoxifen administration (1.3 vs 2.3) and at 2 weeks upon tamoxifen (1.4 vs 5.5) administration, in Krt14-CREER/YFP and Krt14-CREER/SmoM2 respectively.

To test whether tumor cells have distinct morphological features and guided by our theoretical results, we performed two types of quantifications. Firstly, taking 2D cross-sections across tumor and wild-type clones, we calculated the “wetting” angle between lateral membranes at the clonal boundary and the BM as outlined in Fig5c-d. The average was 90° for wild-type cells, as expected because lateral membranes need on average to be vertical with respect with the BM (Fig5e). In tumor however, we found that the contact angles are significantly lower (p-values of 0.035 and 0.003 respectively), with a decrease compared to wt of 4° for week-1 tumors and 7° for week-2. Using these contact angles to estimate (more details in (Fig S2 e) the tension ratio *R* between wt and SmoM2-expressing cells predicted *R* ∼ 1.05 and ∼ 1.10 for week 1 and week 2 respectively.

Secondly, to verify these results independently, we used a different quantification strategy based on the basal area *A_b_* of wild-type and tumor cells. To compute it, we calculated the area of a clone (GFP-positive) that was in contact with the basal lamina (stained using B4-integrin positive) using Imaris software (see Methods for more details), and divided it by the number of basal cells in the clone (Fig. 5d). To correct for the fact that tumor cells have smaller volumes than wild-type cells, we rescaled this basal area by cellular volume 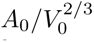, calculated from 3D reconstruction based on GFP labeling of the different clones using Imaris, mirroring the dimensionless ratio calculated in our simulations (Fig. 5). We restrained the analysis to clones consisting of 1 or 2 basal cells, to prevent biases due to larger tumor clone sizes. This revealed that individual SmoM2-expressing cells were more basally attached than wild-type cells at 1w, by around 15%, in qualitative agreement with the model. At 2w, the relatively small number of tumor clones of 1-2 cells resulted in non significant differences. More quantitatively and taking into account standard error of the means in the measurements, this translated into a predicted tension ratio *R* = 1 − 1.2, broadly consistent with the first quantification method.

**Figure 5.**
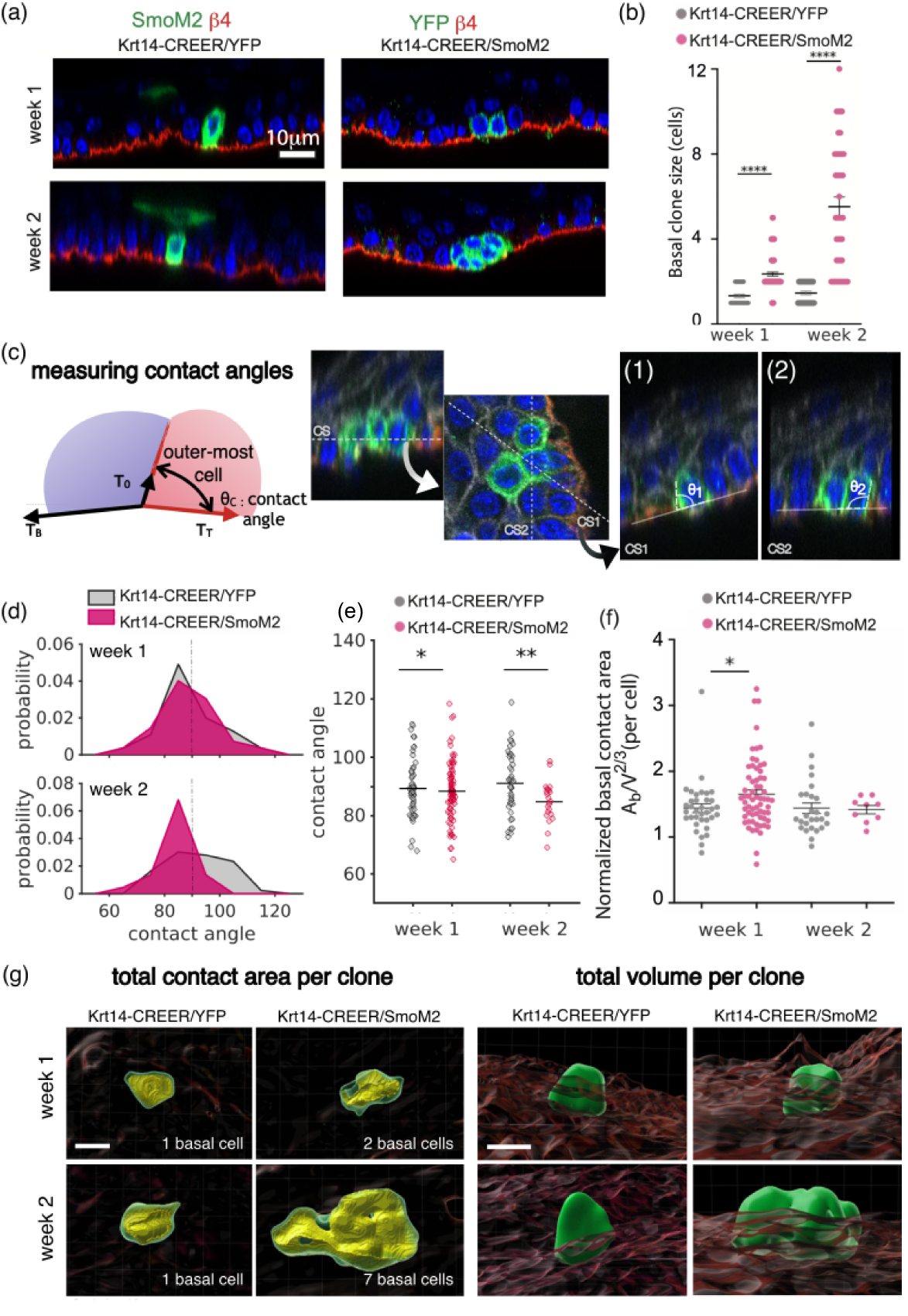
Experimental test of the predicted morphometric signatures of tumor cells. (a)) Immunostaining for *β*4-integrin, YFP and SmoM2 in Krt14-CREER/YFP and Krt14-CREER/SmoM2 epidermis at different time points upon tamoxifen administration. (b) Quantification of the basal clone size in clones from Krt14-CREER/YFP and Krt14-CREER/SmoM2 at different time points upon tamoxifen administration. (c) Schematic depicts the quantity that is used to measure tension homogeneity between adjacent clones– contact angle (*θ_c_*). For more details see Sec:Methods. Using representative z-stack images of mouse skin (IFE), showing Phallodin (gray), DAPi (blue) for nuclei, immunostaining for *β*4-integrin (red) for BM and SmoM2 (green), an example for contact angle measurement is shown for a 3-cell clone. More details in Sec:Methods. (d-e) Contact angle distributions (d) and averages (e) are shown for week 1 (top) and week 2 (bottom). Dotted gray line is drawn at 90°for reference. With time Krt14-CREER/YFP clones spreads more and across 90°, while Krt14-CREER/SmoM2 clones do the opposite.(f) Basal contact area of singlet and doublet clones, normalised by cellular volumes shown for Krt14-CREER/YFP and Krt14-CREER/SmoM2 clones. For week 1 clones, there is a roungh 15% increase in the normalised contact area. The difference between week-2 distributions is not statistically significant due to a smaller number of singlet and doublet clones.(g) Representative images of mouse skin (IFE) is shown for Krt14-CREER/YFP and Krt14-CREER/SmoM2 clones showing the 3D reconstruction of the BM (*β*4-integrin) in brown and clones in green, for times-1 and 2 weeks after tamoxifen administration. On the left panel yellow highlights, the contact area with BM that is the overlapping region between both the 3D objects. Scale bars: 10 µm.

Importantly, a tension ratio of around *R* = 1.05 predicted from our simulations a clonal imbalance of around Δ = 40%, which is comparable in magnitude to clonal imbalances of 25% − 30% that we had found upon Smoothened activation in Involucrin-expressing progenitors or K14-expressing stem cells.

## Discussion

Using a 3D vertex-based model, with proliferation restricted to the bottom-most layer of a multilayered tissue due to niche factors, we show that mechanics and competition for finite space can organically give rise to homeostasis via balanced stochastic fate choices. Indeed, overcrowding in the basal layer is mechanically unstable, which leads to initially symmetric divisions being resolved into asymmetric fate outcomes at the population level. Thus, without any explicit fate assignment to the progenies, we can recover salient aspects found lineage tracing experiments and stochastic models in the absence of mechanics, such as effectively stochastic fate choices and neutral drift of clones at the population level. This generically predicts a distribution of 1*/*4 − 1*/*2 − 1*/*4 for basal-basal, basal-suprabasal and suprabasal-suprabasal fate outcomes, consistent in observations in several systems such as mouse tail or ear epidermis [43, 55]. Interestingly, other organs display rarer symmetric divisions (e.g. 10 − 15% instead of the predicted 25%) [13, 41], which is a signature of more complex regulation that could be added to our model in the future. One simple assumption that could be easily relaxed is the one that divisions take place in the basal plane: if divisions are also allowed to be perpendicular to the BM, this will clearly bias the fraction of asymmetric divisions via purely geometrical effects. Another, more complex, more of regulation would be to consider more explicitly fate choice heterogeneity in the basal layer, for instance via signalling interactions between two daughter cells to regulate their respective fates [31].

Furthermore, beyond clone size evolution, our numerical simulations also makes a number of predictions on how the mechanical state of the tissue affects not only cellular shape index (as studied over recent years), but also clonal shape and dispersion, generalizing previous findings in 2D monolayers linking tissue fluidity to clonal dispersion [7, 11]. In adult skin, clones are typically highly cohesive [55], indicative of a solid tissue rheology, something which would be tested more quantitatively in the future by mechanical measurements.

Surprisingly, upon introduction of mechanical inhomogeneity in the form of lowered basal tension for a sub-population of cells, we discover that the homeostatic balance rapidly becomes tilted towards symmetric divisions. For instance, as little as 10% of disparity in basal tension is enough to induce tumor-like invasiveness within the basal layer, without any change in division rate. Given previous findings which have connected the impaired differentiation, rather than enhanced proliferation, of basal cells in BCC progression, this hinted at the idea that differential mechanical changes could drive differential fate and invasion in tumors. One advantage of our mechanical model is that its core constituent is cellular shape, meaning that tension differences between WT and tumor cells manifest themselves in predictable differential shapes between the two populations. In particular, both the contact angle of lateral surfaces with the BM and the fraction of cellular area in contact with the BM depend predictably on the tension differences, facilitating the inference of this critical parameter in the data.

We then proceeded to measure detailed cellular morphometrics in both wild-type and tumor cells at different time points, and found that both metrics were consistent in pointing towards a tension imbalance in tumor cells of 5 − 10%, which is sufficient in the model to reproduce the magnitude of imbalance towards symmetric renewal observed in long-term lineage tracing experiments [55].

Thus, this model underlines how mechanical forces can be key to tissue dynamics not only as regulators of tissue shape, but also by modulating cellular fate choices in settings where proliferative potential depends on tissue location. Although we have concentrated here on the skin epidermis, our model is highly general and could apply to any stratified tissue. In addition to such mechanical effects, a logical next step would be to consider more direct feedback of forces through fate via mechano-sensing [1, 45, 58], as well as considering more complex tissues with more heterogeneous fates [46]. Another interesting aspect of differential mechanics would be to consider the influence of spatial modulation of ECM mechanics, which has recently been shown to critically modulate the competence of tumor initiation in skin [2]. Finally, although we have concentrated on confluent tissues of fixed densities, wound healing from basal cell ablation has been shown to be linked to changes in tissue fluidity [52]. Overall, this works shows that mechanical models can also be a powerful tool for stem cell biology, by connecting easily accessible morphological signatures such as cell shape to fate and tumor invasiveness in the initial stages of cancer.

## Methods

### Mechanical parameters in the simulation

We begin with the simplest homeostatic model where every cell has the same preferred cell shape index *s*_0_ irrespective of where it belongs in the three-layered tissue. While we also explore for a fluid-like shape of *s*_0_ = 5.5 (Fig panels: S1, S2 and S3) to find both-homeostatic and tumor behavior comparable to the solid shape of *s*_0_ = 5.3, we stick to the later as it resembles the physiological tissue better, in particular due to the absence of significant clonal fragmentation in experiments (Fig S3). These cells are contained between two almost-flat surfaces with a fixed base-line tension value of *T_B_* = 1.0 (for *s*_0_ = 5.5, the value is much smaller of 0.1). These value are chosen based on the region of the shape-tension phase space with no tension-induced pinning [49]. We augment this further by zooming into the tension range and making sure to have no artificial pinning behaviour, as shown in in (Fig S1 e-f). The cells also have the same noise level throughout the tissue set by self-propulsion speed of *v*_0_ = 0.1, which is not a special value as the dynamical behavior is highly robust with respect to noise (Fig S1 g-h) till a threshold value above which the basement membrane fails to stabilise into a flat surface.

### Details about noise implementation

To test the effect of noise, we add cellular activity on top of the static model. As we use Voronoi cells in our system, there is one degree of freedom per cell, which is the Voronoi center. The equation of motion is implemented in a straight-forward way [49]. Mechanical force on a cell is computed using the derivative of the non-dimensionalised energy functional (e) from Eq:2 i.e. **f***_i_* = *∂e/∂***r***_i_*. Cellular activity, with respect to the cellular surroundings (such as extracellular matrix for instance), is then added such that it has a polarization vector **n̂***_i_* and active force due to self-propulsion of *v*_0_*/µ* where *v*_0_ is the propulsion speed, and *µ* is the cell mobility (it is set to unity).

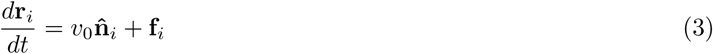

The polarization vector evolves via white Gaussian noise on a unit sphere with diffusion coefficient *D_r_* as shown below-

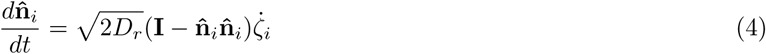

where **I** is a 3×3 identity tensor, **n̂***_i_***n̂***_i_* is the dyadic product of the polarization vector with itself, and *ζ_i_* is the white Gaussian noise.

### Timescales in the simulations

A natural time unit in the simulations is 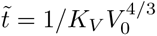 which we get by non-dimensionalising the dynamical equations, and thus is set to unity. For implementing noise, we use a self-propulsion speed of *v*_0_ = 0.1 and rotational noise *D_r_* = 1, essentially transitioning to a Brownian regime as quickly as *t > t̃/D_r_*. The characteristic diffusion timescale in a homogeneous three-dimensional system with *s*_0_ = 5.3 is *τ*_0_ = 10^5^*t̃* (for *s*_0_ = 5.5 it is 10^3^*t̃*). Note that we do not focus here on the effect of self-propulsive activity and persistence, which have been extensively explored [5], given our focus in this system is on the role of cellular division and mechanics-dependent fate choices. We use an integration time-step of Δ = 0.01*t̃*.

Cell divisions happen exclusively in the basal layer, that is defined as all the cells which have their centers within a one cell length distance from the BM and have a finite connection to the BM. In this work, the division timescale of a given basal cell is on an average - *τ_D_* ∼ 3900*t̃*, making the division rate *k_D_* = 0.00026*t̃^−^*^1^. This frequency is chosen in order to have less division-induced-fluidization of the tissue (to resemble a physiological tissue) and at the same time be computationally feasible i.e. a full simulation completes under 24 hours. It has been observed that for a 2D confluent monolayer in the solid regime, the chosen division rate diffuses cells only by a couple of cell lengths by the end of our simulation time [11].

An individual cell division event occurs in every 50*t̃*. A randomly chosen mother cell is allowed to grow to a preferred cell volume of 3.5 for 25*t̃*. Note that this value of preferred volume is larger than the volume actually reached by cells due to the properties of the regular Voronoi tessalation, which is the reason we use a relatively larger value of 3.5. In the future, different tessalation methods such as Radical Voronoi could be used to dampen this effect. The preferred cell surface area is modified accordingly to maintain the preferred shape index. After this, the mother cell is replaced by two daughter cells with centers of mass at the same height as the mother (i.e. in-plane division) and placed 0.25 unit lengths (a unit length refers to the typical cell length *l*_0_) from the mother cell’s position in a co-linear fashion. This axis of division in the plane of the BM is constructed using a randomly chosen angle with respect to the X axis, and is bisected by the division plane. The daughters inherit all the mechanical attributes and the clonal group number of the mother cell. The system is evolved for another 25*t̃* to adjust to the new configuration. After this cycle of 50 timesteps, another mother cell is chosen at random, and the process is sequentially repeated.

The time it takes for the system to reach steady-state comprises of two stages-in the first stage, the system is evolved for *T_SS_*_1_ = 1000*t̃* with no cellular divisions so that the basement membrane stabilises to a flat surface [49]. In the second stage, we run simulations with divisions for *T_SS_*_2_ = 4000*t̃* to ensure at least all basal cells have divided once, and to let the system come to a steady state with respect to proliferation. By this time, the basal density therefore achieves a steady state as shown in (Fig S1 c-d). Only after this stage, every basal cell is allotted a clonal group number, and for the tumor case, inhomogeneities in basal tensions are implemented for four cells equidistant from each other. The simulation is then run for a total of 11000*t̃* that is ∼ 2.8*τ_D_*. The linear fitting of clonal growth dynamics is analysed for the last ∼ 1*τ_D_* window, which is when the homeostatic clones achieve a steady state, as one can observe in 2. The transition from linear to exponential is captured really well with Eq:5, which we use to complement the fate distribution findings. The definition of Δ that we use for fate distributions, is slightly different from this, since it is easier to understand in terms of population asymmetry as opposed to a ratio– Δ −→ 2*r*Δ̃. [22]

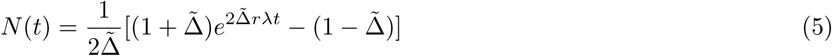

As open-boundaries in Voronoi models are highly non-trivial to implement, we allow three more layers comprising of *pseudo* cells below the basement membrane, that sandwich the tissue via periodic boundary conditions along z-axis. Since their only job is to sustain the basal surface, these cells are only active during the SS1 stage (with the same diffusion profile as real cells) and are mechanically pinned for the entirety of the simulation. They will have negligible feedback to the real cells across BM because we use low basal tension values that are much below the tension-induced-pinning regime which is when the cells across interface become coupled [49].

### Analysis protocol for cell-fate distribution in simulations

Lineage trees for every clone is plotted using the mother-daughter cell information. Every cell has a unique ID, and we construct lineage information between mothers and daughters at each division event. Using this information, we construct lineage trees with respect to time as shown in Fig4b. At the end of the simulation, for every lineage tree we have the list of all the clonal members both– past (eliminated through top layer) and present (basal and non-basal). For the clones that survive till the end we also have the list of all the final basal members. To identify the set of members that have extruded we simply remove the set of final basal members from the set of all members. This way we also identify the daughters that continued to divide or have survived till the end (nodes are shown in pink) and the ones that extruded (nodes shown in blue). The daughters that survive till the final timepoint of the simulation, are not included in fate distribution because they need more time for their fate to become apparent (i.e. either extrude or divide again).

Next, we use this information for finding the probabilities of each fate outcome as a function of simulation parameters. For each cell division event, we check how many of the daughter cells have extruded, and we measure the statistics of each of the three possible fate outcomes, i.e. none (basal-basal division), one (basal-suprabasal), or both (suprabasal-suprabasal) of the cells have extruded. We then filter out the divisions that occur after one complete turnover i.e *t* ≥ 1*τ_D_*, because not all the basal cells have gotten the chance to differentiate, thereby artificially biasing the distribution towards ‘symmetric basal’ fate. From the remaining division events, we compute the proportion for each fate, for the given value of tension ratio *R*, as plotted in Fig4.

### Quantitative morphometric analysis of clone and cell shapes

We next quantify morphological features of the simulations at both-single-cell and clone level.

As for the quantification of single cell basal area *A_T_*, we use the final snapshot i.e. 11000*t̃*, to quantify it. This includes cells from every kind of clone including the singlets. We isolate all the facets that are in contact with the BM and sum it up.

With an increase in noise, we observe that the solid clonal shapes in solid parameter regime becomes less compact and sprawl more. For fluid systems we see that while the tumor clones are compact in zero noise, they undergo explicit fragmentation for higher noise levels (Fig S3).

To infer tension inhomogeneity between adjacent clones in a manner that can be mapped to experimental measurements, we use the contact angle (*θ_c_*) that their lateral contact forms with the BM. In simulations we typically use a tumor consisting of a single cell,so shorter ∼ 1*τ_D_*-simulations are also used to identify the lateral edges and their coordinates. As the basement membrane is on an average parallel to the XY plane, we compute the angles made by the edges with the XY plane. This is plotted for different tumors in Fig S2 e.

To measure the same in experimental images, Figure panel Fig5 c-d explains the entire protocol with a schematic and example. For a clone with a relatively flat BM, an outer-basal-cell is chosen (depicted in pink in Fig5c) to measure the angle between its lateral edges and the BM. Figure Fig5d shows an example for contact angle measurement for a 3-cell clone shown using orthogonal view on the top-left. Using the apical view on right (Z-stack corresponding to CS), several cross-sectional cuts are made, two of which are shown for this case - CS1 and CS2. Angles-*θ*_1_ and *θ*_2_ are measured between the outer-most lateral edges and the BM line (solid gray line) for cross-sections CS1 and CS2 in sub-figures-(1) and (2).

### Mice

Krt14-CreER,Rosa/YFP and Rosa/SmoM2-YFP mice were obtained from the JAX repository. Mouse colonies were maintained in a certified animal facility in accordance with European guidelines for the laboratory animal use and care based on the 2010/63/EU Directive. Experiments involving mice presented in this work were approved by the Animal Welfare and Ethics Body, Direção-Geral da Alimentação e Veterinária (DGAV,Portuguese Authority) under protocol DGAV protocol number 011681.

### Skin tumour induction

For tumour induction 1.5-months-old mice were used. Krt14-CREER/Rosa-SmoM2 and received an intraperitoneal injection of 0.1 mg (0.5 mg/ml) of tamoxifen (ref. T5648-0005, Sigma). Mice were sacrificed and analysed at different time points following tamoxifen administration. Krt14-CREER/Rosa-SmoM2 and Krt14-CREER/Rosa-YFP were heterozygous for the Rosa-YFP and Rosa-SmoM2 mutation in the Rosa Locus. The tail of these animals was used in our analysis.

### Lineage tracing experiments during skin homeostasis

For the lineage tracing experiments 1.5-months-old mice were used. Krt14-CREER/Rosa-YFP mice received an intraperitoneal injection of 0.1 mg (0.5 mg/ml) of tamoxifen (ref. T5648-0005, Sigma) Mice were sacrificed and analysed at different time points following tamoxifen administration. All mice were heterozygous for Rosa-YFP.

### Whole-mounts of tail epidermis

Whole mounts of tail epidermis were performed as previously described [55]. Specifically, pieces of tail were incubated for 1 hour (h) at 37 °C in EDTA 20 mM in PBS in a rocking plate, then using forceps the dermis and epidermis were separated and the epidermis was fixed for 30 min in 4% Formaldehyde methanol-free (ref FB002, ThermoFisher Scientific) in agitation at room temperature and washed 3 x with PBS. For the immunostaining, tail skin pieces were blocked with blocking buffer for 3 h (PBS, horse serum 5%, Triton 0.8%) in a rocking plate at room temperature. After, the skin pieces were incubated with primary antibodies diluted in blocking buffer overnight at 4 °C, the next day they were washed with PBS-Tween 0.2% for 3 × 10 min at room temperature, and then incubated with the secondary antibodies diluted in blocking buffer for 3 h at room temperature, washed 2 × 10 min with PBS-Tween 0.2% and washed for 10 min in PBS. Finally, they were incubated in Hoechst (1:1000) diluted in PBS for 30 min at room temperature in the rocking plate, washed 3 × 10 min in PBS and mounted in DAKO mounting medium supplemented with 2.5% Dabco (Sigma). Primary antibodies used were the following: Goat anti-GFP (1:800, ref. ab6673, Abcam), Rat anti-*β*4-integrin (1:500, ref. 553745, BD Pharmingen). *R*&*D* Systems Secondary antibodies used were the following: anti-goat, AlexaFluor488 (Goat, ref A-11055, Invitrogen). Samples were also stained with Phalloidin conjugated to AlexaFluor647 (1:400, ref A30107, ThermoFisher Scientific).Images were acquired using Z-stacks with an inverted confocal microscope LSM 980 (Carl Zeiss).

### Analysis of clone size

The quantification of the the basal and total clone size were determined by counting the number of SmoM2-positive cells (in Krt14-CREER/Rosa-SmoM2 mice) and YFP-postive cells (in Krt14-CREER/Rosa-YFP mice) in each clone using orthogonal views of the whole-mount tail epidermis in the interscale region of interfollicular epidermis as described in [55]. *β*4-integrin staining was used to classify the clone according to their location in basal or suprabasal layers.

### Image analysis

The “Surface” module that is part of the Imaris 9.6 software (Andor Inc.) was used to carry out the 3D-rendering of the clone structure labelled with anti-GFP and the basal membrane structure labelled with anti-*β*4-integrin. The 3D-rendering of the objects was performed using smoothing 1 *µm*, background subtraction 12 µm and similar thresholding settings across all experiments for the morphological analysis of the clone volume (CV) and the clone area (CA). Structures outside of the rendered objects were excluded from final analysis. The proximity between the basal cell surface object and the basal membrane object were determined using “Surface Contact Area” (SCA) Imaris XTension integrated plugin (https://imaris.oxinst.com/open/view/surface-surface-contact-area). The contact area between basal cells from the clone and the basal membrane was calculate using the following formula: 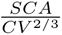.

### Statistical analysis

All statistical analysis were performed using GraphPad Prims v.8.0.1 software. Data are expressed as mean ± s.e.m. Normality was tested using Shapiro-Wilk Test. For the database that followed normality, Pvalues were estimated with unpaired t-test for two experimental groups or for multiple comparison two-way ANOVA. For the dataset that did not follow the normal distribution, Pvalues were calculated with the Mann-Whitney test. For all the figures, the number of mice and the number of clones is indicated in the figure legend.

## Author contribution

Conceptualization: P.S., A.S-D., E.H. Simulations: P.S, Experiments: S. C., R.S.; Supervision: A.S-D., E.H., Manuscript writing: P.S., E.H., A.S-D with inputs from all authors

## Acknowledgments

We thank Alois Schlögl, Paula Sanematsu, Susana Moreno Flores, Bernat Corominas-Murtra, Stefania Tavano, Gayathri Singharaju, and Hannezo group members for helpful discussions, the Bioimaging facilty at ISTA as well as Matthias Merkel and Lisa Manning for sharing the 3D Voronoi code. We also thank the Champalimaud animal facility, and Anna Pezzarossa and Champalimaud ABBE platform for the help with microscopy and image processing. This work was supported by EMBO (ALTF 522-2021), a Fundação para a Cîencia e Tecnologia grant to A.S.D (PTDC/MED-ONC/5553/2020), as well as the European Research Council (grant 851288 to EH). A.S.D, S.C and R.M.S is supported by QuantOCancer Project Horizon European Union’s Horizon 2020 programme (grant agreement No 810653).

## Supplementary Figures

**Figure S1.**
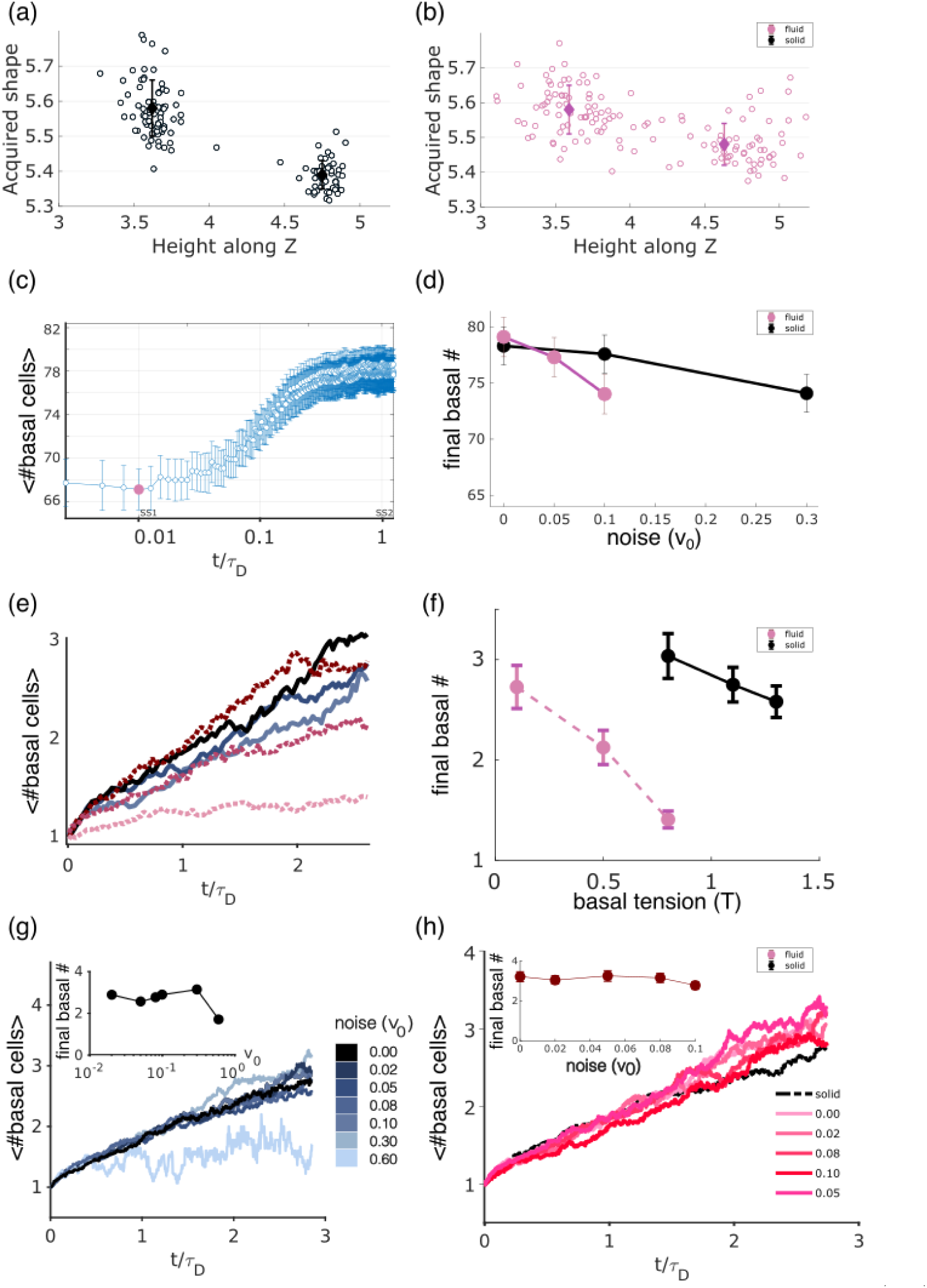
Cellular morphometrics and sensitivity analysis in homeostatic simulations. (a-b) Shape index of cells along the Z-axis for simulations in the solid regime (a, black, shape index=5.3) and fluid regime (b, pink, shape index=5.5), showing stronger shape differences in basal cells for solid tissues. (c) Number of basal cells as a function of time in simulations. At early times (before time indicated as SS1 in pink), tissues are equilibrated without divisions. After this time, basal cells divisions are allowed, and the total basal cell number increased to reach an homeostatic steady-state plateau. (d) Steady-state value of basal cell number as a function of noise activity *v*_0_, and tissue rheology (black is solid and in pink is fluid). e) Temporal evolution and f) Sensitivity analysis of the Average basal clone size as a function of basal tension for solid or fluid tissues. g-h) Temporal evolution of the average basal clone size for solid rheology (g) and fluid rheology (h) for different values of noise activity *v*_0_. Insets show the robustness wrt noise.

**Figure S2.**
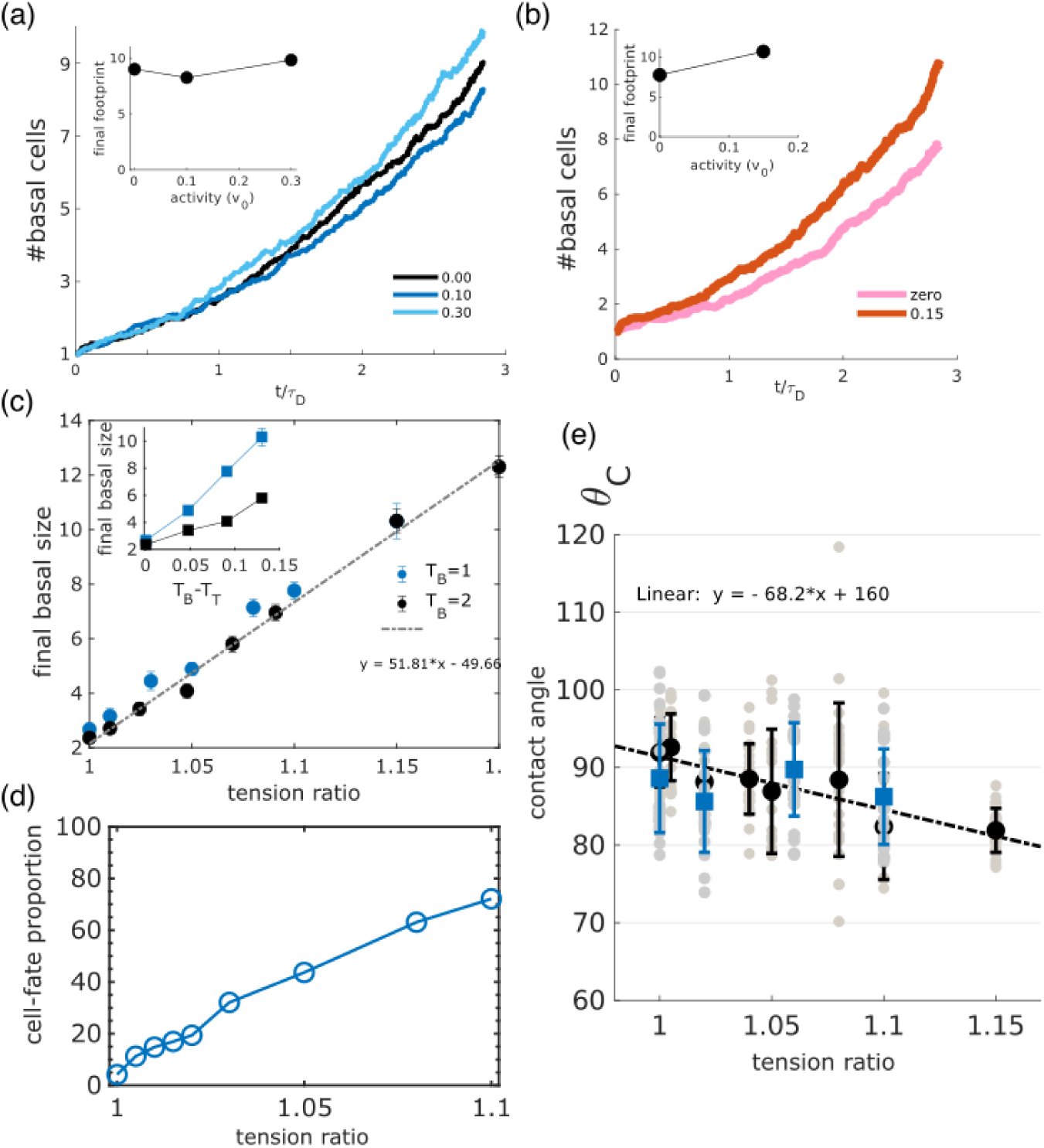
Cellular morphometrics and sensitivity analysis in tumor simulations. (a-b) Temporal evolution for the average basal clone size of tumors for a solid (a, shape index=5.3) and fluid (b, shape index=5.5) rheology. Different colors indicate in each panel different values for the noise activity *v*_0_. Insets show the final clone size as a function of noise activity, showing only a weak dependence. c) Final average basal clone size as function of the tension imbalance ratio *R* (or absolute tension imbalance difference *T_B_* − *T_T_*, inset) for tumor clones, for different values of the basement tension (*T_B_*=1 in blue, *T_B_* = 2 in black). Normalizing tensions (tension ratio) provides a good prediction between final basal size and *R*, even for different absolute values of basement tension *T_B_*, with the relationship being well-described by a linear fit (dashed line). d) Predictions from the simulations for the relationship between fate imbalance Δ and tension ratio, showing a strong correlation. (e) Contact angles computed in simulations, plotted against R. For both *T_B_* = 1.0 (grey) and *T_B_* = 2.0 (orange), we see a linear decline with respect to R, which is independent of the absolute values of tension as seen in panel c. The dotted curve is the linear fitting to the normal base-line tension i.e. *T_B_* = 1.0.

**Figure S3.**
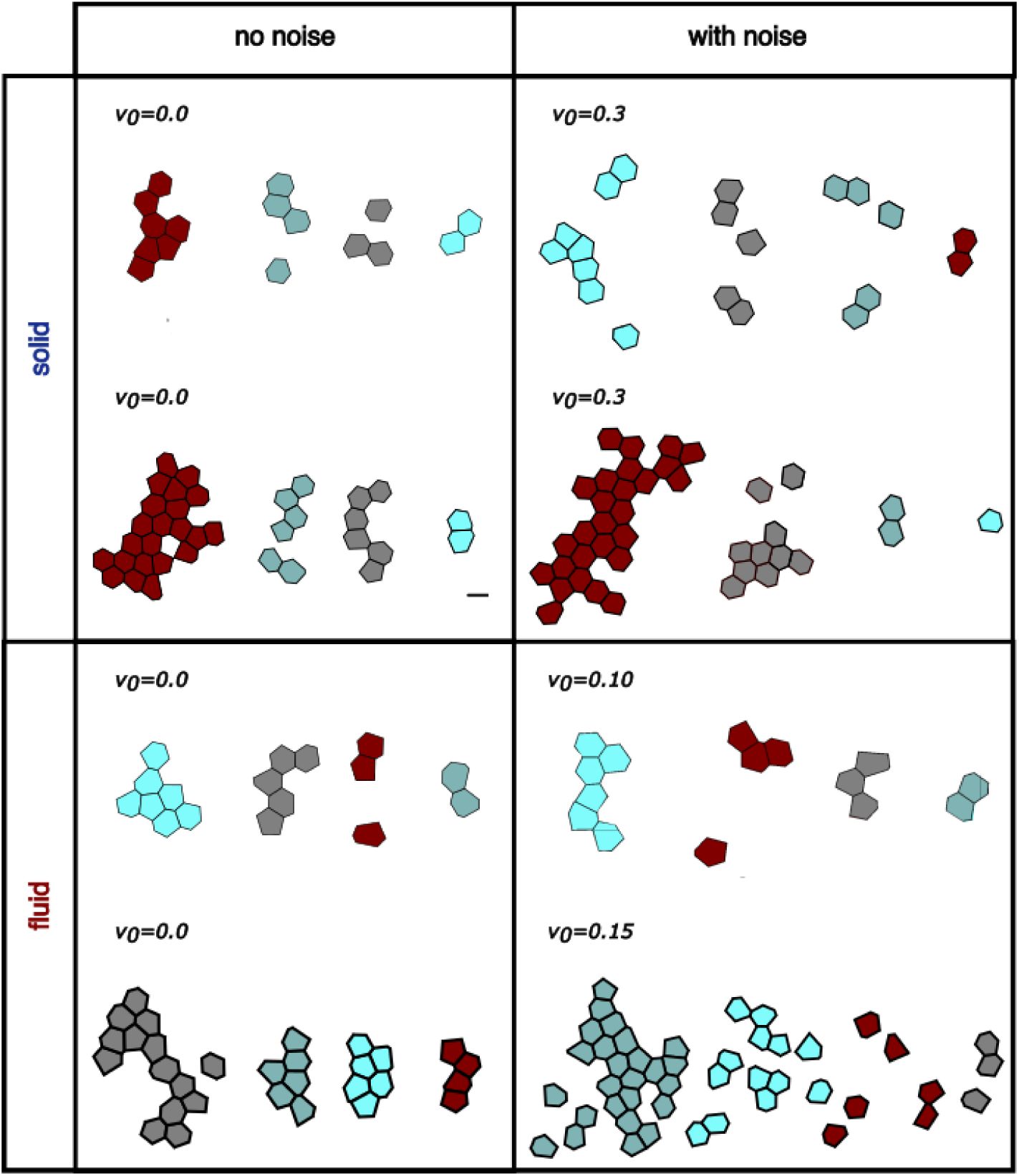
Clonal shape for different model parameters. We display for each set of parameters four simulated clones, showing both differences in single cell and clonal shape morphometrics. Left column indicates no noise activity (*v*_0_ = 0), right column indicates finite noise activity). Top line indicates solid rheology (shape index= 5.3), bottom line indicates fluid rheology (shape index = 5.5). In each panel, top clones indicate homeostatic simulations while bottom clones indicate tumor simulations (*R* = 1.1). As expected, the effect of noise activity is to create clonal dispersion and irregular clone shapes.

**Figure S4.**
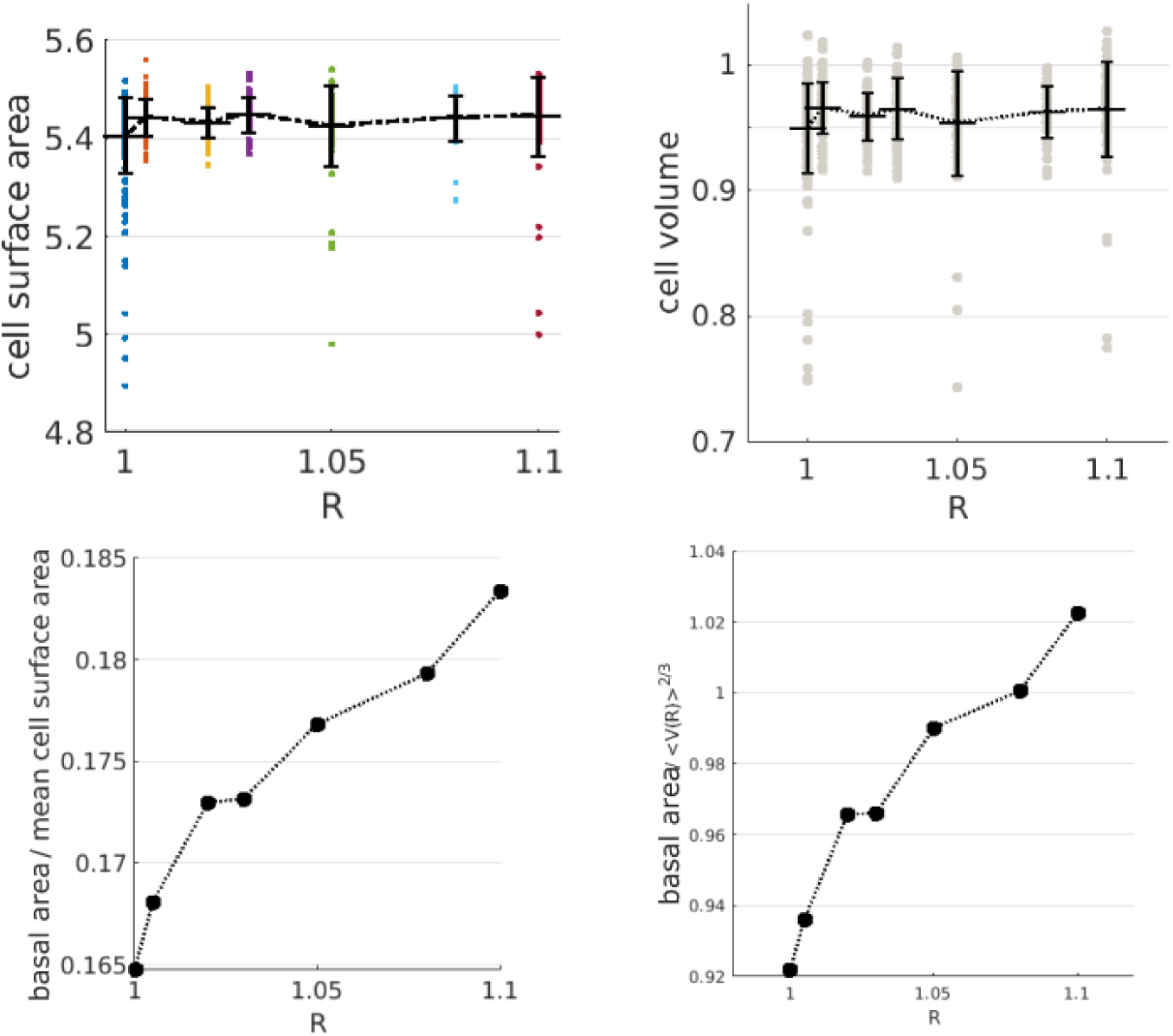
Cell and clonal morphometrics in tumor simulations. a-b) Total basal cell area (a) and volumes (b) are nearly unaffected by changes in the basal tension ratio *R* in simulations. c-d) Cellular areas in contact with the BM (normalized either with respect to total cell area in panel c, or cell volume *V* ^2^*^/^*^3^ in panel d) shows a monotonic increase with tension ratio *R*, due to the energetic preference of tumor cells to adhere to the BM.

## References

1. M. Aragona, A. Sifrim, M. Malfait, Y. Song, J. V. Herck, S. Dekoninck, S. Gargouri, G. Lapouge, B. Swedlund, C. Dubois, P. Baatsen, K. Vints, S. Han, F. Tissir, T. Voet, B. D. Simons, and C. Blanpain. Mechanisms of stretch-mediated skin expansion at single-cell resolution. Nature 2020 584:7820, 584:268–273, 7 2020.

2. N. Bansaccal, P. Vieugue, R. Sarate, Y. Song, E. Minguijon, Y. A. Miroshnikova, D. Zeuschner, A. Collin, J. Allard, D. Engelman, A. L. Delaunois, M. Liagre, L. de Groote, E. Timmerman, D. V. Haver, F. Impens, I. Salmon, S. A. Wickström, A. Sifrim, and C. Blanpain. The extracellular matrix dictates regional competence for tumour initiation. Nature, 623:828–835, 11 2023.

3. N. Barker. Adult intestinal stem cells: critical drivers of epithelial homeostasis and regeneration. Nature reviews. Molecular cell biology, 15:19–33, 1 2014.

4. D. Bi, J. H. Lopez, J. M. Schwarz, and M. L. Manning. A density-independent rigidity transition in biological tissues. Nat. Phys., 11:1074–1079, 12 2015.

5. D. Bi, X. Yang, M. C. Marchetti, and M. L. Manning. Motility-driven glass and jamming transitions in biological tissues. Phys. Rev. X, 6:021011, 2016.

6. C. Blanpain and B. D. Simons. Unravelling stem cell dynamics by lineage tracing. Nature reviews. Molecular cell biology, 14:489–502, 8 2013.

7. L. Bocanegra-Moreno, A. Singh, E. Hannezo, M. Zagorski, and A. Kicheva. Cell cycle dynamics control fluidity of the developing mouse neuroepithelium. Nature Physics, 19, 2023.

8. K. K. Chiou, L. Hufnagel, and B. I. Shraiman. Mechanical stress inference for two dimensional cell arrays. PLOS Computational Biology, 8:e1002512, 2012.

9. B. Corominas-Murtra and E. Hannezo. Modelling the dynamics of mammalian gut homeostasis. Seminars in Cell & Developmental Biology, 150-151:58–65, 12 2023.

10. B. Corominas-Murtra, C. L. Scheele, K. Kishi, S. I. Ellenbroek, B. D. Simons, J. V. Rheenen, and E. Hannezo. Stem cell lineage survival as a noisy competition for niche access. Proceedings of the National Academy of Sciences of the United States of America, 117:16969–16975, 7 2020.

11. M. Czajkowski, D. M. Sussman, M. C. Marchetti, and M. L. Manning. Glassy dynamics in models of confluent tissue with mitosis and apoptosis. Soft Matter, 15, 2019.

12. D. P. Doupé, M. P. Alcolea, A. Roshan, G. Zhang, A. M. Klein, B. D. Simons, and P. H. Jones. A single progenitor population switches behavior to maintain and repair esophageal epithelium. Science, 337:1091–1093, 8 2012.

13. D. P. Doupé, A. M. Klein, B. D. Simons, and P. H. Jones. The ordered architecture of murine ear epidermis is maintained by progenitor cells with random fate. Developmental Cell, 18:317–323, 2 2010.

14. G. Erdemci-Tandogan, M. J. Clark, J. D. Amack, and M. L. Manning. Tissue flow induces cell shape changes during organogenesis. Biophysical Journal, 115:2259–2270, 12 2018.

15. R. Farhadifar, J. C. Röper, B. Aigouy, S. Eaton, and F. Jülicher. The influence of cell mechanics, cell-cell interactions, and proliferation on epithelial packing. Current Biology, 17:2095–2104, 12 2007.

16. V. F. Fiore, M. Krajnc, F. G. Quiroz, J. Levorse, H. A. Pasolli, S. Y. Shvartsman, and E. Fuchs. Mechanics of a multilayer epithelium instruct tumour architecture and function. Nature, 585:433, 9 2020.

17. E. Hannezo, J. Prost, and J. F. Joanny. Theory of epithelial sheet morphology in three dimensions. Proceedings of the National Academy of Sciences of the United States of America, 111:27–32, 1 2014.

18. Y. C. Hsu and E. Fuchs. Building and maintaining the skin. Cold Spring Harbor Perspectives in Biology, 14:a040840, 7 2022.

19. D. J. Jörg, Y. Kitadate, S. Yoshida, and B. D. Simons. Stem cell populations as self-renewing many-particle systems. 10.1146/annurev-conmatphys-041720-125707, 12:135–153, 3 2021.

20. S. Kim, M. Pochitaloff, G. A. Stooke-Vaughan, and O. Campàs. Embryonic tissues as active foams. Nature Physics 2021 17:7, 17:859–866, 4 2021.

21. Y. Kitadate, D. J. Jörg, M. Tokue, A. Maruyama, R. Ichikawa, S. Tsuchiya, E. Segi-Nishida, T. Nakagawa, A. Uchida, C. Kimura-Yoshida, S. Mizuno, F. Sugiyama, T. Azami, M. Ema, C. Noda, S. Kobayashi, I. Matsuo, Y. Kanai, T. Nagasawa, Y. Sugimoto, S. Takahashi, B. D. Simons, and S. Yoshida. Competition for mitogens regulates spermatogenic stem cell homeostasis in an open niche. Cell Stem Cell, 24:79–92.e6, 1 2019.

22. A. M. Klein, D. E. Brash, P. H. Jones, and B. D. Simons. Stochastic fate of p53-mutant epidermal progenitor cells is tilted toward proliferation by uv b during preneoplasia. Proceedings of the National Academy of Sciences of the United States of America, 107:270–275, 2010.

23. A. M. Klein, D. P. Doupé, P. H. Jones, and B. D. Simons. Mechanism of murine epidermal maintenance: Cell division and the voter model. Physical Review E - Statistical, Nonlinear, and Soft Matter Physics, 77:031907, 3 2008.

24. A. M. Klein and B. D. Simons. Universal patterns of stem cell fate in cycling adult tissues. Development, 138:3103–3111, 8 2011.

25. P. L. Krapivsky, S. Redner, and E. Ben-Naim. Spin Dynamics. Cambridge University Press, 11 2011.

26. A. D. Lander, K. K. Gokoffski, F. Y. Wan, Q. Nie, and A. L. Calof. Cell lineages and the logic of proliferative control. PLOS Biology, 7:e1000015, 1 2009.

27. I. C. Mackenzie. Relationship between mitosis and the ordered structure of the stratum corneum in mouse epidermis. Nature 1970 226:5246, 226:653–655, 1970.

28. C. Malinverno, S. Corallino, F. Giavazzi, M. Bergert, Q. Li, M. Leoni, A. Disanza, E. Frittoli, A. Oldani, E. Martini, T. Lendenmann, G. Deflorian, G. V. Beznoussenko, D. Poulikakos, K. H. Ong, M. Uroz, X. Trepat, D. Parazzoli, P. Maiuri, W. Yu, A. Ferrari, R. Cerbino, and G. Scita. Endocytic reawakening of motility in jammed epithelia. Nature Materials 2017 16:5, 16:587–596, 1 2017.

29. G. Mascŕe, S. Dekoninck, B. Drogat, K. K. Youssef, S. Brohée, P. A. Sotiropoulou, B. D. Simons, and C. Blanpain. Distinct contribution of stem and progenitor cells to epidermal maintenance. Nature, 489(7415):257—262, September 2012.

30. M. Merkel and M. L. Manning. A geometrically controlled rigidity transition in a model for confluent 3d tissues. New J. Phys., 20:022002, 2 2018.

31. K. R. Mesa, K. Kawaguchi, K. Cockburn, D. Gonzalez, J. Boucher, T. Xin, A. M. Klein, and V. Greco. Homeostatic epidermal stem cell self-renewal is driven by local differentiation. Cell Stem Cell, 23, 2018.

32. Y. A. Miroshnikova, H. Q. Le, D. Schneider, T. Thalheim, M. Rübsam, N. Bremicker, J. Polleux, N. Kamprad, M. Tarantola, I. Wang, M. Balland, C. M. Niessen, J. Galle, and S. A. Wickström. Adhesion forces and cortical tension couple cell proliferation and differentiation to drive epidermal stratification. Nature cell biology, 20:69–80, 1 2018.

33. J. A. Mitchel, A. Das, M. J. O’Sullivan, I. T. Stancil, S. J. DeCamp, S. Koehler, O. H. Ocaña, J. P. Butler, J. J. Fredberg, M. A. Nieto, D. Bi, and J. A. Park. In primary airway epithelial cells, the unjamming transition is distinct from the epithelial-to-mesenchymal transition. Nature Communications 2020 11:1, 11:1–14, 10 2020.

34. A. Mongera, P. Rowghanian, H. J. Gustafson, E. Shelton, D. A. Kealhofer, E. K. Carn, F. Serwane, A. A. Lucio, J. Giammona, and O. Campàs. A fluid-to-solid jamming transition underlies vertebrate body axis elongation. Nature 2018 561:7723, 561:401–405, 9 2018.

35. T. Nagai and H. Honda. A dynamic cell model for the formation of epithelial tissues. Philosophical Magazine B, 81:699–719, 2001.

36. C. G. Neumann. The expansion of an area of skin by progressive distention of a subcutaneous balloon: Use of the method for securing skin for subtotal reconstruction of the ear. Plast. Reconstruct. Surg., 19:124–130, 1957.

37. C. M. Niessen, M. L. Manning, and S. A. Wickström. Mechanochemical principles of epidermal tissue dynamics. Cold Spring Harbor Perspectives in Biology, page a041518, 6 2024.

38. W. Ning, A. Muroyama, H. Li, and T. Lechler. Differentiated daughter cells regulate stem cell proliferation and fate through intra-tissue tension. Cell stem cell, 28:436, 3 2021.

39. T. Otani, T. Ichii, S. Aono, and M. Takeichi. Cdc42 gef tuba regulates the junctional configuration of simple epithelial cells. The Journal of cell biology, 175:135–146, 10 2006.

40. J. A. Park, J. H. Kim, D. Bi, J. A. Mitchel, N. T. Qazvini, K. Tantisira, C. Y. Park, M. McGill, S. H. Kim, B. Gweon, J. Notbohm, R. Steward, S. Burger, S. H. Randell, A. T. Kho, D. T. Tambe, C. Hardin, S. A. Shore, E. Israel, D. A. Weitz, D. J. Tschumperlin, E. P. Henske, S. T. Weiss, M. L. Manning, J. P. Butler, J. M. Drazen, and J. J. Fredberg. Unjamming and cell shape in the asthmatic airway epithelium. Nature Materials, 14:1040–1048, 10 2015.

41. G. Piedrafita, V. Kostiou, A. Wabik, B. Colom, D. Fernandez-Antoran, A. Herms, K. Murai, B. A. Hall, and P. H. Jones. A single-progenitor model as the unifying paradigm of epidermal and esophageal epithelial maintenance in mice. Nature Communications 2020 11:1, 11:1–15, 3 2020.

42. D. Pinheiro, R. Kardos, Édouard Hannezo, and C. P. Heisenberg. Morphogen gradient orchestrates pattern-preserving tissue morphogenesis via motility-driven unjamming. Nature Physics 2022 18:12, 18:1482–1493, 10 2022.

43. P. Rompolas, K. R. Mesa, K. Kawaguchi, S. Park, D. Gonzalez, S. Brown, J. Boucher, A. M. Klein, and V. Greco. Spatiotemporal coordination of stem cell commitment during epidermal homeostasis. Science (New York, N.Y.), 352:1471–1474, 6 2016.

44. A. Roshan, K. Murai, J. Fowler, B. D. Simons, V. Nikolaidou-Neokosmidou, and P. H. Jones. Human keratinocytes have two interconvertible modes of proliferation. Nature cell biology, 18:145–156, 1 2016.

45. M. Rübsam and C. M. Niessen. Stretch exercises for stem cells expand the skin. Nature 2021 584:7820, 584:196–198, 7 2020.

46. M. Rübsam, R. Püllen, F. Tellkamp, A. Bianco, M. Peskoller, W. Bloch, K. J. Green, R. Merkel, B. Hoffmann, S. A. Wickström, and C. M. Niessen. Polarity signaling balances epithelial contractility and mechanical resistance. Scientific Reports, 13, 2023.

47. P. Sahu. Fluidization and segregation in confluent models for biological tissues. Dissertations - ALL, 8 2020.

48. P. Sahu, J. Kang, G. Erdemci-Tandogan, and M. L. Manning. Linear and nonlinear mechanical responses can be quite different in models for biological tissues. Soft Matter, 16:1850–1856, 2020.

49. P. Sahu, J. M. Schwarz, and M. L. Manning. Geometric signatures of tissue surface tension in a three-dimensional model of confluent tissue. New Journal of Physics, 23:093043, 9 2021.

50. P. Sahu, D. M. Sussman, M. Rübsam, A. F. Mertz, V. Horsley, E. R. Dufresne, C. M. Niessen, M. C. Marchetti, M. L. Manning, and J. M. Schwarz. Small-scale demixing in confluent biological tissues. Soft Matter, 16:3325–3337, 4 2020.

51. P. C. Sanematsu, G. Erdemci-Tandogan, H. Patel, E. M. Retzlaff, J. D. Amack, and M. L. Manning. 3d viscoelastic drag forces contribute to cell shape changes during organogenesis in the zebrafish embryo. Cells & Development, 168:203718, 12 2021.

52. R. M. Sarate, J. Hochstetter, M. Valet, A. Hallou, Y. Song, N. Bansaccal, M. Ligare, M. Aragona, D. Engelman, A. Bauduin, O. Campàs, B. D. Simons, and C. Blanpain. Dynamic regulation of tissue fluidity controls skin repair during wound healing. Cell, 187:5298–5315.e19, 9 2024.

53. D. B. Staple, R. Farhadifar, J.-C. Röper, B. Aigouy, S. Eaton, and F. Jülicher. Mechanics and remodelling of cell packings in epithelia. Eur. Phys. J. E, 33:117–127, 2010.

54. D. M. Sussman, J. M. Schwarz, M. C. Marchetti, and M. L. Manning. Soft yet sharp interfaces in a vertex model of confluent tissue. Physical Review Letters, 120, 1 2018.

55. A. Śanchez-Dańes, E. Hannezo, J. C. Larsimont, M. Liagre, K. K. Youssef, B. D. Simons, and C. Blanpain. Defining the clonal dynamics leading to mouse skin tumour initiation. Nature, 536:298–303, 7 2016.

56. C. Villeneuve, A. Hashmi, I. Ylivinkka, E. Lawson-Keister, Y. A. Miroshnikova, C. Pérez-Gonźalez, S. M. Myllymäki, F. Bertillot, B. Yadav, T. Zhang, D. M. Vignjevic, M. L. Mikkola, M. L. Manning, and S. A. Wickström. Mechanical forces across compartments coordinate cell shape and fate transitions to generate tissue architecture. Nature Cell Biology 2024 26:2, 26:207–218, 2 2024.

57. A. Yanagida, E. Corujo-Simon, C. K. Revell, P. Sahu, G. G. Stirparo, I. M. Aspalter, A. K. Winkel, R. Peters, H. D. Belly, D. A. Cassani, S. Achouri, R. Blumenfeld, K. Franze, E. Hannezo, E. K. Paluch, J. Nichols, and K. J. Chalut. Cell surface fluctuations regulate early embryonic lineage sorting. Cell, 185:777–793.e20, 3 2022.

58. A. M. Zöllner, M. A. Holland, K. S. Honda, A. K. Gosain, and E. Kuhl. Growth on demand: Reviewing the mechanobiology of stretched skin. Journal of the Mechanical Behavior of Biomedical Materials, 28:495–509, 12 2013.

